# Galanin receptor 1 expressing neurons in hippocampal-prefrontal circuitry modulate goal directed attention and impulse control

**DOI:** 10.1101/2024.07.29.605653

**Authors:** Fany Messanvi, Vladimir Visocky, Carolyn Senneca, Kathleen Berkun, Maansi Taori, Sean P. Bradley, Huikun Wang, Hugo Tejeda, Yogita Chudasama

## Abstract

Neuropeptides like galanin are increasingly recognized as modulators of cognitive pathways. Galanin has been implicated in a wide range of pathological conditions in which frontal and temporal structures are compromised. Recently, we discovered that direct pharmacological stimulation of galanin receptor type 1 (GalR1) in the ventral prefrontal cortex (vPFC) and ventral hippocampus (vHC) caused opposing effects on attention and impulse control behaviors. In the present study, we investigate how neurons expressing GalR1 in these two areas differentially contribute to these behaviors. First, using multiplex fluorescent in situ-hybridization, we established that GalR1 is predominantly expressed in glutamatergic neurons in both the vPFC and vHC. Rats were assessed in their visuospatial attention and impulse control behaviors using the 5- Choice task. We developed a novel viral approach to gain genetic access to GalR1-expressing neurons in the vPFC and vHC, and found that optogenetic excitation of GalR1 expressing neurons in the vPFC, but not vHC, selectively disrupted attention. Finally, using fiber photometry, we measured the bulk calcium dynamics in GalR1-expressing neurons and discovered that GalR1- expressing neurons in the vPFC and vHC showed opposing activity; increased activity in neurons in the vPFC corresponded to correct, attentive actions, whereas activity in the vHC signaled impulsive errors. This region- and response-specific intrinsic activity of galanin, mediated by subclasses of neurons in frontotemporal circuitry participates in shaping the expression of executive-control behaviors that often go awry in various disorders of mental health.

## Introduction

Cognitive deficits in attention and response control manifest as behavioral symptoms of distraction and impulsivity which ultimately dysregulate normal executive functioning. These symptoms underlie a range of pathological conditions marked by a lack of self-control contributing to episodes of violence, deviant sexual behavior, and pathological gambling [1]. They also constitute the main difficulty in neurodevelopmental disorders characterized by limited focus, hyperactivity and repetitive behaviors [2]. Such behaviors are thought to be mediated by abnormal activity in the prefrontal cortex and other brain areas known to be involved in the normal control of attention and fine-tuned by ascending modulatory systems [3–7]. The catecholamine noradrenaline (NA), which originates in the locus coeruleus (LC), is a neuromodulator best known for regulating prefrontal-cognitive functions under high arousing conditions [8]. Extracellular NA has pro-cognitive effects by enhancing attentional mechanisms and controlling impulsive urges [9–11]. Recent work has suggested that these behaviors are controlled by specific projections emanating from the LC to the dorsal and ventral divisions of the prefrontal cortex [12], but it is unclear if the neuromodulatory effect of NA is specific to fine-tuning of activity in the prefrontal cortex alone, or optimized with other neuronal circuitries involved in cognition.

In addition to NA, LC neurons co-express several neuropeptides, particularly galanin which is found in eighty percent of those neurons in the rat [13]. This co-existence and the presence of galanin receptors in regions such as the prefrontal cortex and hippocampus [14, 15] strongly implicate this neuropeptide in the noradrenergic modulation of cognitive control processes. Studies that link galanin and cognition relate mostly to its role in learning and memory functions and its potential involvement in Alzheimer’s disease pathophysiology [16]. The relationship between galanin and noradrenaline with respect to attentional control has not been systematically explored. In mice, for example, galanin overexpression has little impact on attentional performance [17], but these studies lacked methods that enabled both neuroanatomical and receptor specificity. At the cellular level, galanin inhibits the activity of LC neurons in vitro [18, 19] and enhances NA-induced inhibition of LC neurons [20]. In the cerebral cortex, galanin decreases the NA-induced cyclic AMP response [21]. Since galanin has no detectable action when applied alone, both NA and galanin must work together for efficient noradrenergic transmission [21]. Moreover, although galanin is released when galanin expressing neurons fire at high frequency [22–24], the behavioral conditions contributing to galanin release have not been identified.

Recently, we discovered that galanin, through local stimulation of galanin receptor type 1 (GalR1), affects cognitive control functions in rats through its direct actions in the ventral prefrontal cortex (vPFC) and the ventral hippocampus (vHC) [25]. The main change to behavior concerned the rate of impulsive premature responding. In the vPFC, this stimulation led to a high rate of impulsive responses, whereas in the vHC it had the opposite effect, making rats more controlled in their responses and therefore more successful. Notably, high impulsivity led to poor control of visual attention suggesting that the actions of GalR1 in the vHC and vPFC facilitate the normal control of behavior.

In the present study we use multiple approaches to characterize the functional differences between GalR1-expressing neurons of the vPFC and vHC and their involvement in complex cognitive behavior. We assessed behavior using the 5-Choice task, a well-established test of executive function in rats modeled after its human analogue, the continuous performance test. We genetically targeted the neurons expressing GalR1 and captured the rapid dynamic properties of these neurons in the vPFC and vHC using fiber photometry. Since local stimulation of GalR1 in the vPFC and vHC produce opposing behavioral effects [25], we surmised that GalR1-expressing neurons in the vHC and vPFC differentially signal cognitive mechanisms of attention and impulse control that shape the executive response.

## Results

### Galanin receptor 1 is expressed in glutamatergic cells in the vPFC and the vHC

To better understand the region-specific mechanisms of GalR1 actions, we started by investigating whether galanergic markers were differentially expressed and distributed in the vPFC and the vHC. We first examined the presence of galanin fibers in both regions of interest. Galanin- immunoreactive fibers and terminals were present along the entire dorsoventral extent of the PFC (Fig. S1a-d). In the HC, the density of galanin fibers was consistent across the dentate gyrus and CA1-CA3 fields (Fig. S1e-h). Next, we characterized the distribution of the GalR1 mRNA. *In situ* hybridization using RNAscope confirmed the presence of GalR1 mRNA in both vPFC and vHC subregions. Within the PFC, the highest density of GalR1 mRNA was in the IL cortex (Fig. 1a-c) located preferentially in layer 5 (Fig. 1c). In the vHC, the distribution was greatest in the pyramidal layers of the vCA1 and ventral subiculum (vSub) (Fig. 1d-f). These observations were largely consistent with previous reports [14, 15, 26]. We then determined the cell-type distribution of GalR1 mRNA in the IL cortex and vCA1/vSub since both areas showed the highest expression of GalR1 mRNA (Figs. 1g, and l). In both cases, the majority of GalR1 mRNA was expressed in glutamatergic neurons (Figs. 1h-i and m-n; upper panels). A much smaller proportion of the GalR1 was expressed in GABAergic neurons (Figs. 1j-k and o-p; lower panels), reflecting the lower abundance of this class of neurons. Together, these results indicate that vPFC and vHC circuits can be modulated by GalR1 actions upon neurons residing in specific layers and subregions.

**Figure 1.**
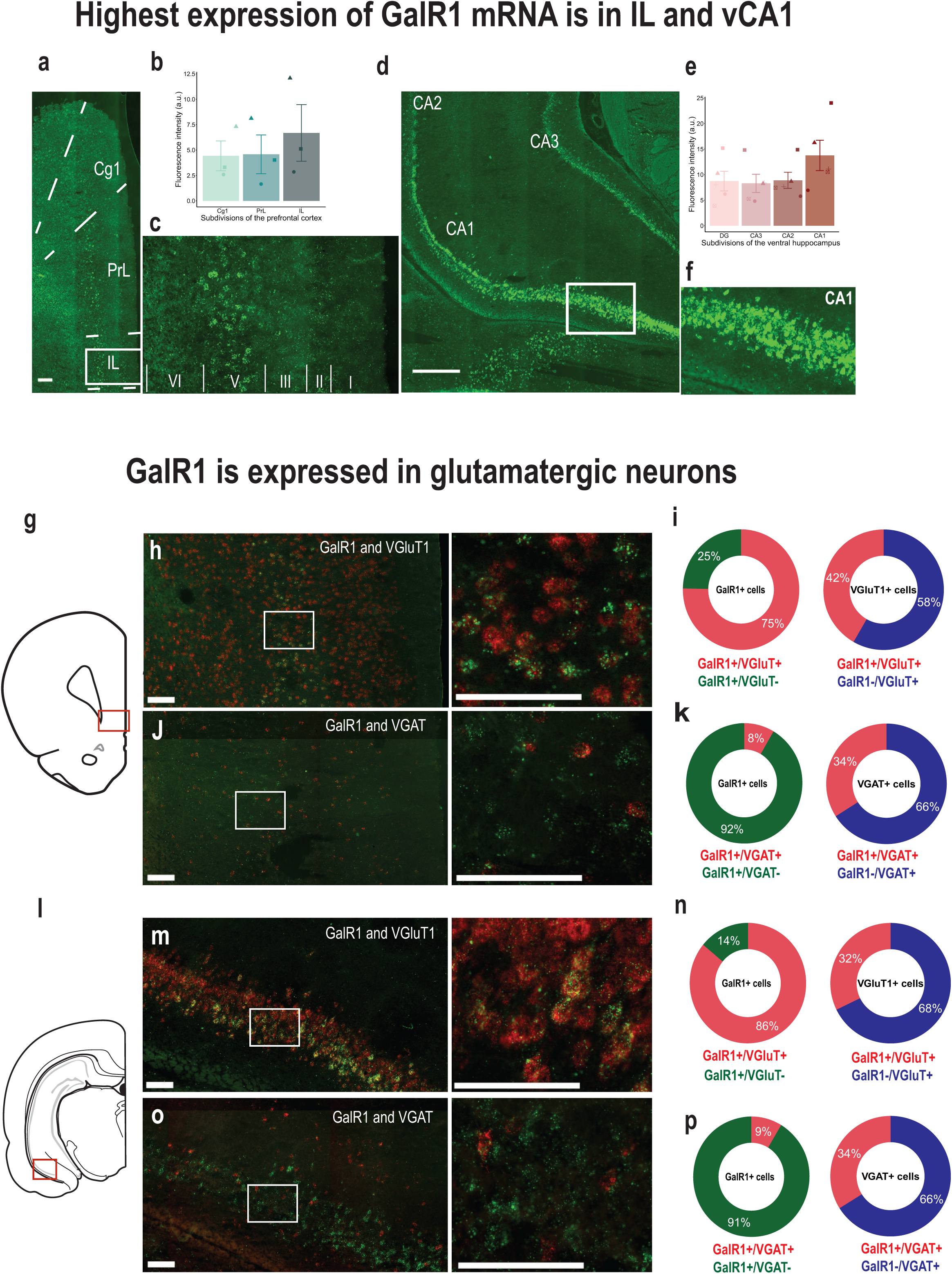
Expression of GalR1 in the vPFC and the vHC. **(a)** Representative image of the distribution of GalR1 in the PFC (scale bar: 200 µm). **(b)** Quantification of GalR1 mRNA fluorescence intensity in three subdivisions of the PFC (N = 3 animals, 3-4 sections per animal; Cg1: 4.43 ± 1.47, PrL: 4.58 ± 1.91, IL: 6.69 ± 2.79, arbitrary unit: au). Bar chart represents mean ± SEM. Dots represent individual animals. **(c)** Magnified image of the distribution of GalR1 in different cortical layers of the IL cortex. **(d)** Representative image of the distribution of GalR1 in the vHC (scale bar: 500 µm). **(e)** Quantification of GalR1 mRNA fluorescence intensity in the vHC (N = 5 animals, 3-4 sections per animal; DG: 8.73 ± 1.92, CA3: 8.29 ± 1.78, CA2: 8.89 ± 1.58, CA1: 13.76 ±2.97, arbitrary unit: au). Bar chart represents mean ± SEM. Dots represent individual animals. **(f)** Magnified image of the distribution of GalR1 in the different layers of the vCA1/vSub. **(g-k)** Co-expression of GalR1 mRNA with glutamatergic neuron marker VGluT1 (GalR1+/VGluT1+ from 642 GalR1+ cells: 75.2 ± 3.2 %, GalR1+/VGluT1- from 642 GalR1 cells: 24.6 ± 3.3 %; GalR1+/VGluT1+ from 229 VGluT1+ cells: 58.3 ± 5.2 %, GalR1-/VGluT1+ from 229 VGluT1+ cells: 41.7 ± 5.2 %; 3 animals, 3-4 sections per animal) and the GABAergic neuron marker VGAT (GalR1+/VGAT+ from 647 GalR1+ cells: 8.4 ± 1.0 %, GalR1+/VGAT- from 647 GalR1+ cells: 91.6 ± 1.0 %; GalR1+/VGAT+ from 83 VGAT+ cells: 66.0 ± 7.4 %, GalR1-/VGAT+ from 83 VGAT+ cells: 34.0 ± 7.4 %; 2 animals, 2-4 sections per animal) in the IL. Scale bars: 100 µm. **(l-p)** Co-expression of GalR1 mRNA with glutamatergic neuron marker VGluT1 (GalR1+/VGluT1+ from 590 GalR1+ cells: 86.0 ± 0.9 %, GalR1+/VGluT1- from 590 GalR1+ cell: 14.0 ± 0.9 %; GalR1+/VGluT1+ from 340 VGluT1+ cells: 67.8 ± 3.4 %, GalR1-/VGluT1+ from 340 VGluT1+ cells: 32.2 ± 3.4 %; 3 animals, 2 sections per animal) and the GABAergic neuron marker VGAT (GalR1+/VGAT+ from 610 GalR1+ cells: 8.6 ± 1.3 %, GalR1+/VGAT- from 610 GalR1+ cells: 91.4 ± 1.3 %; GalR1+/VGAT+ from 74 VGAT+ cells: 66.1 ± 1.3 %, GalR1-/VGAT+ from 74 VGAT+ cells: 33.9 ± 1.3 %; 1 animal, 3 sections) in the vCA1. Scale bars: 100 µm.

### Selective stimulation of GalR1-expressing neurons affects behavioral performance

To better understand the causal relationship between the neurons expressing GalR1 in these brain regions and attentional control of behavior, we selectively stimulated the activity of these neurons via temporally targeted optogenetic techniques. A genetic construct expressing Cre recombinase under the control of GalR1 promoter was packaged into a AAV1 to target the GalR1- expressing neurons (Fig. 2a). We first validated the construct by quantifying its specificity in the vPFC (Fig. 2b, c) and verified the absence of fluorescence in the cerebellum which has low GalR1 expression (Fig. 2d). We then injected the GalR1-Cre virus into the vHC and vPFC, using different fluorophore reporters in the two areas, allowing us to determine the distinct projections of their GalR1 expressing neurons (Fig. 2e). We found that vPFC and vHC GalR1-expressing neurons project widely to brain areas involved in attention and impulse control including the midline thalamus, ventral striatum and septum (Fig. 2f-i)). Interestingly, GalR1-expressing neurons in the vHC selectively targeted the deep layers of the vPFC running amid the cell bodies of GalR1- expressing neurons (Fig. 2f).

**Figure 2.**
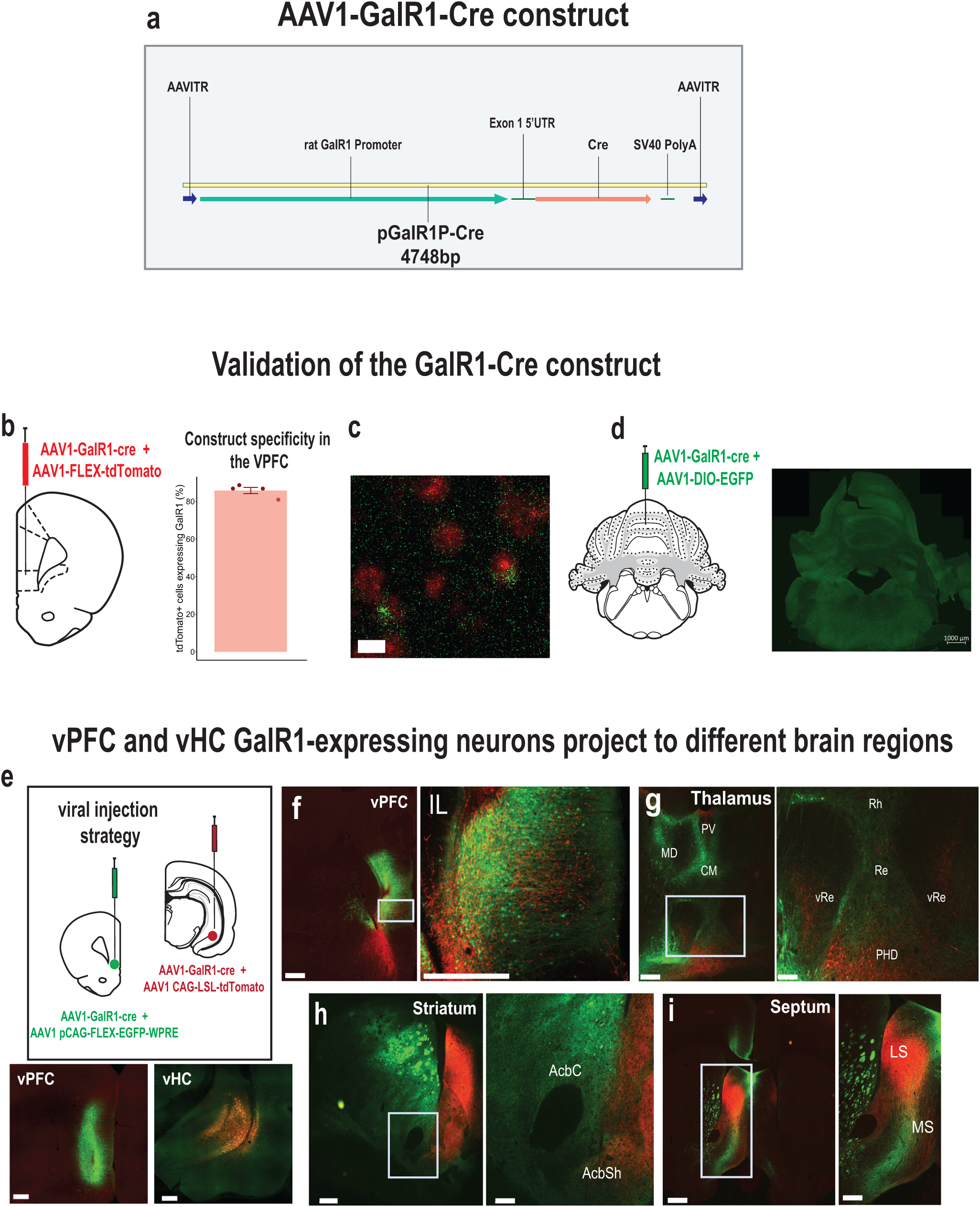
Validation of the GalR1-Cre construct. **(a)** Schematic of AAV vector construct expressing Cre recombinase under the control of the GalR1 promoter. **(b)** Injection strategy and assessment of the specificity of the virus in the vPFC. Quantification of the cells expressing the Cre-dependent fluorophore that were also positive for GalR1 (85.8 ± 1.7 %; 771 cells, N = 4 animals, 3-6 sections per animal). Bar chart represents mean ± SEM. Dots represent individual animals. (c) Representative image showing co-expression of GalR1 mRNA in green and tdTomato in cells infected with the viral construct **(d)** Injection strategy to validate the specificity of the GalR1-Cre construct in the cerebellum (negative control) with corresponding microphotograph showing no expression **(e)** viral strategy shows GalR1-Cre virus injected together with the reporter protein in the vPFC (green) and vHC (red) (upper panel). Representative photomicrographs showing injection sites in the vPFC (left) and the vHC (right) (lower panel, scale bar: 500 µm). **(f)** Dense presence of vPFC GalR1-expressing neurons within the infralimbic cortex, and vHC GalR1 fibers near the vPFC injection site. vHC fibers are concentrated to the deep layers. (scale bars: 500 µm and 100 µm) **(g)** vPFC and vHC GalR1 neurons project to distinct parts of the thalamus. The Rhomboid (Rh) and Reuniens (Re) nuclei receive dense vPFC projections but scarce vHC projections (scale bars: 500 µm and 200 µm). **(h)** vPFC and vHC GalR1 neurons project to distinct parts of the striatum. In the nucleus accumbens, vPFC projections are seen in the core and shell, while vHC fibers are mostly seen in the shell (scale bars: 500 µm and 200 µm). **(i)** vPFC and vHC GalR1 neurons project to the lateral septum with GalR1 fibers from both regions targeting the dorsal and intermediate divisions (scale bars: 500 µm and 200 µm). IL, infralimbic; MD, mediodorsal; PV, paraventricular; CM, centromedial; Rh, rhomboid nucleus; Re, nucleus reuniens; vRe, ventral nucleus reuniens; PHD, posterior hypothalamic area, dorsal part; AcbC, accumbens core; AcbSh, accumbens shell; LS, lateral septum; MS, medial septum.

To investigate the functional contribution of these neurons, we examined the behavioral effects of selectively exciting GalR1-expressing neurons. We expressed ChR2 in these neural populations in either the vPFC or vHC and implanted an optic fiber above the viral injection site (Fig. 3a). Following post-operative recovery, rats were trained on the 5-choice task until they acquired a baseline level of performance (Fig. 3b; see methods). We then optically activated the GalR1-expressing neurons during the pre-stimulus interval. We did this in an interleaved fashion such that only half of the trials in each session were stimulated (Fig. 3b).

**Figure 3.**
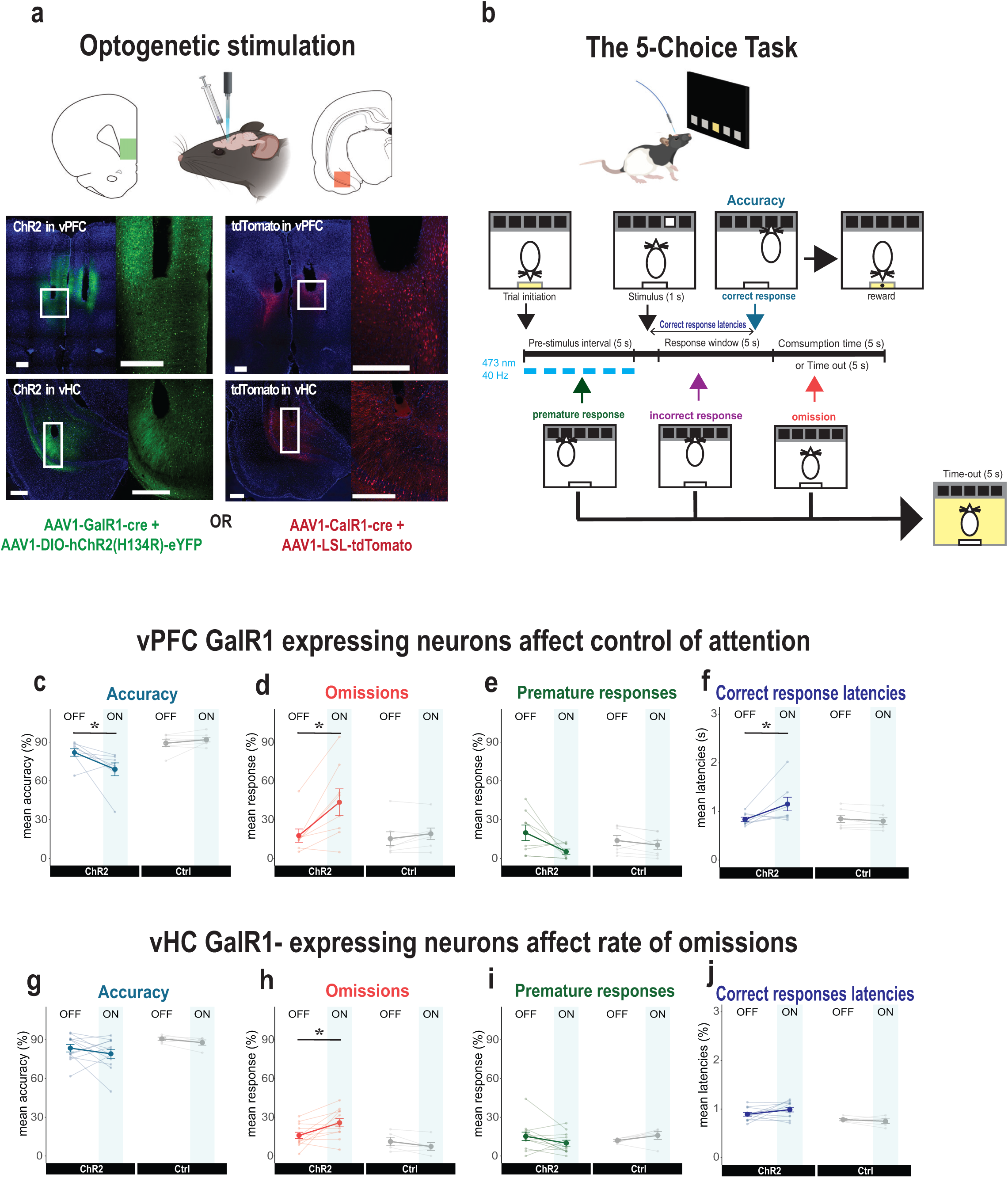
Selective stimulation of GalR1-expressing neurons affects behavioral performance **(a)** Schematic showing virus and fiber optic placement strategy for optical manipulation. Representative images of viral expression in the vPFC (upper panels) and the vHC (lower panels). Scale bars: 500 µm). **(b)** Schematic of the 5-choice task and stimulation protocol. **(c-f)** Effects of optical stimulation of vPFC GalR1-expressing cells on accuracy, omissions, premature responses, and correct response latencies (ChR2: n = 8; Ctrl: n= 7). **(g-j)** Effects of optical stimulation of vHC GalR1-expressing cells on accuracy, omissions, premature responses, and correct response latencies (ChR2: n = 12; Ctrl: n= 5). Error bars represent SEM. * p < 0.05 (pairwise comparison between groups after significant interaction effect in Mixed ANOVA).

Optogenetic activation of GalR1-expressing neurons in the vPFC, affected executive behavior in three ways. First, it reduced the rats’ ability to accurately detect the visual target (Fig. 3c).

Second, it greatly increased the number of trial omissions (Figs. 3d and S4a, b). Third, it increased their latencies to respond correctly (Fig. 3f). It also led to a trend towards a reduction in impulsive responses (Fig. 3e). All other measures including reward collection latency were not impacted (Figs. S2 and S3).

The specificity of the stimulation site in the vPFC was important. While we successfully targeted the ventral infralimbic/prelimbic region in most animals, we noted that some animals had optic fibers implanted rostral to the target site, namely the medial orbital (MO) division of the vPFC (Fig. S5a). These animals showed a different pattern of behavior during optical stimulation compared with the infralimbic/prelimbic division of the vPFC-ChR2 rats (Fig. S5b). Although they were also less accurate in stimulated trials (Fig. S5c), the stimulation did not alter their rate of omissions (Fig. S5d). In addition, they tended to make more premature responses upon photo- stimulation (Fig. S5e). Thus, in agreement with previous reports, optimal performance in cognitive-executive tasks like the 5-choice task require the interaction of different vPFC subdivisions [27, 28].

In contrast to the strong and repeatable effects of exciting GalR1-expressing cells in the vPFC, exciting the same population in the vHC had little impact on performance (Fig. 3g-j). One exception was a significant increase in the number of omissions during stimulation (Figs. 3h, S2, and S4), but all other aspects of behavior were generally intact. Thus, while the GalR1-expressing cell populations of the vPFC have a direct impact on attentional control of behavior, in the vHC these cells potentially impact motivational elements of task performance.

### Activity of GalR1-expressing neurons reflect attention and impulsivity

We next captured the distinct dynamic responses of vPFC and vHC GalR1-expressing neurons during performance of the 5-choice task, using *in vivo* calcium fiber photometry (Fig. 4). The calcium indicator GCamP7f was expressed in GalR1-expressing neurons of the vPFC and the vHC, and an optic fiber was placed above the viral injection site to record changes in fluorescence with a fiber photometry system (Fig. 4a, b). The signals from the vPFC and the vHC were first parsed by trial outcomes and then aligned to different task events within each trial category.

**Figure 4.**
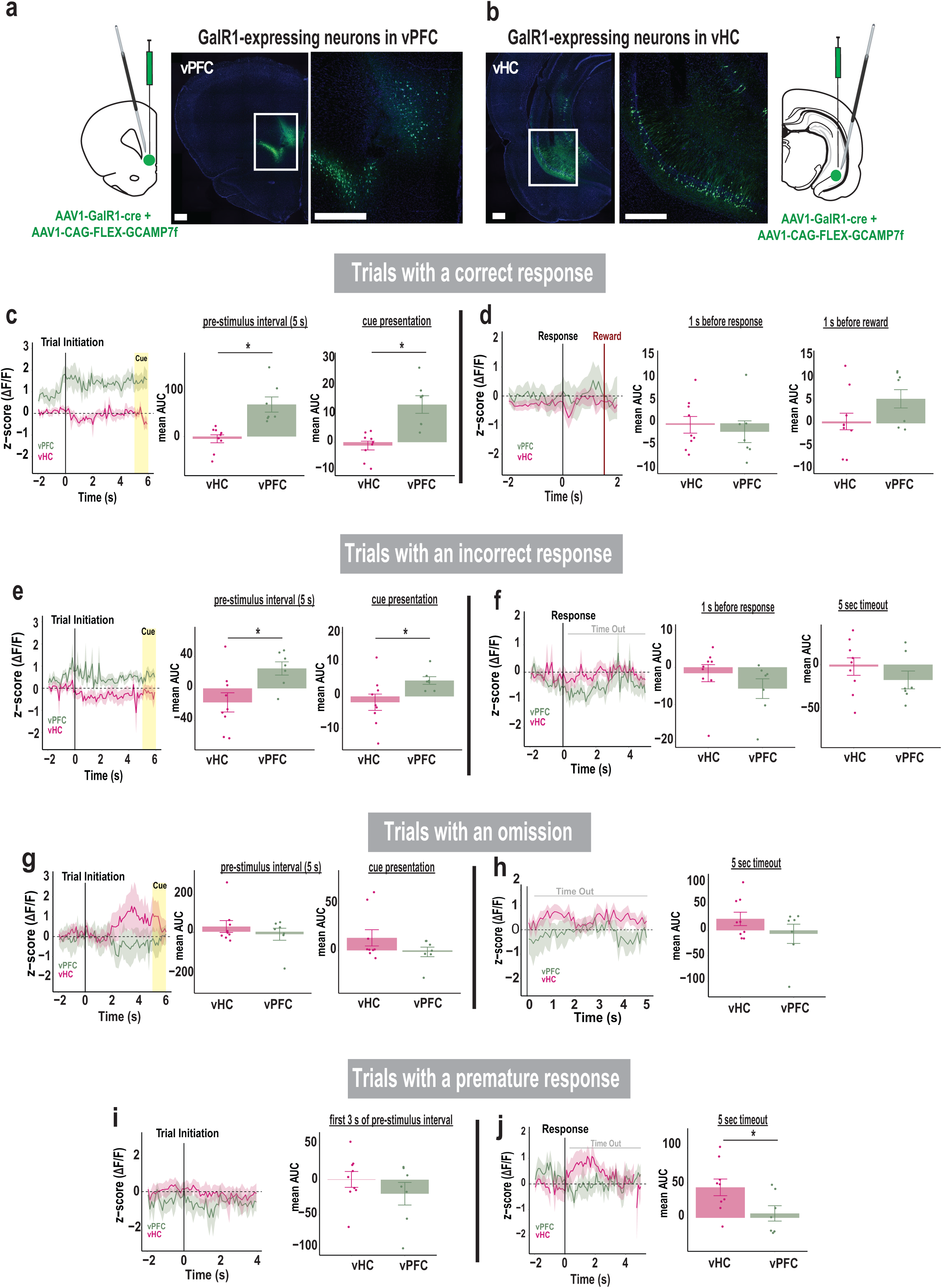
Activity of GalR1-expressing neurons reflects attention and impulsivity **(a)** Schematic of viral injection and optic fiber placement in the vPFC, and **(b)** in vHC, each showing representative image of GCaMP7f expression in each area (scale bars: 500 µm). **(c)** Comparison of calcium signal in vPFC and vHC GalR1-expressing neurons during the trials with a correct response. The signal is aligned to trial initiation. Left: traces represent average activities (mean area under the curve: mean AUC) during the pre-stimulus interval and cue presentation. Right: histograms showing higher vPFC activity compared to vHC during the pre-stimulus interval [t(14) = 4.275, p = 0.001] and the cue presentation [t(14) = 4.472, p = 0.001]. **(d)** Comparison of activity of GalR1-expressing neurons in vPFC and vHC activity during trials with a correct response. Left: traces represent the average activity around the response and reward. The signal is aligned to the response. Right: histograms of vPFC and vHC activity before the response (signal aligned to the response) [t(14) = -0.512, p = 0.616] and before the reward (signal aligned to the reward) [t(14) = 1.652, p = 0.121]. **(e)** Comparison of calcium signal in vPFC and vHC GalR1- expressing neurons during trials with an incorrect response. The signal is aligned to trial initiation. Left: traces represent average activities during the pre-stimulus interval and cue presentation. Right: histograms show average activity of vPFC is higher relative to vHC during the pre-stimulus interval [t(14) = 2.721, p = 0.017] and cue presentation [t(14) = 2.195, p = 0.046]. **(f)** Comparison of vPFC and vHC activity during the trials with an incorrect response. The signal is aligned to the response. Left: traces represent average activity before the response and during the time-out period. Right: histograms show vPFC and vHC activity before the response [t(14) = -1.185, p = 0.256] and during the time-out [t(14) = -1.055, p = 0.309]. **(g)** Comparison of calcium signal in vPFC and vHC GalR1-expressing neurons during the trials with an omission. The signal is aligned to trial initiation. Left: traces represent average activity of neurons during the pre-stimulus interval and cue presentation. Right: histograms show average activity in vPFC and vHC during the pre-stimulus interval [t(14) = -0.904, p = 0.381] and cue presentation [t(14) = -1.380, p = 0.189]. **(h)** Comparison of calcium signal in vPFC and vHC GalR1-expressing neurons during the trials with an omission. The signal is aligned to the end of the response window when an omission is detected. Left: traces represent average activity during the time-out period following an omission. Right: histograms showing the average activity of vPFC and vHC during the time-out period [t(14) = 0.577, p = 0.573]. **(i)** Comparison of calcium signal in vPFC and vHC GalR1-expressing neurons during the trials with a premature response. The signal is aligned to trial initiation. Left: traces represent average activities during the first 3 seconds of the pre-stimulus interval. Right: histograms showing vPFC and vHC average activity during the pre-stimulus interval [t(14) = - 1.064, p = 0.305]. (**j**) Comparison of calcium signal in vPFC and vHC GalR1-expressing neurons during the trials with a premature response. The signal is aligned to the premature response. Left: traces represent average activity before the response and during the time-out period. Right: Average activity of vPFC and vHC during the time-out [t(14) = -2.241, p = 0.042]. (vPFC in green, n = 7; vHC in pink, n= 9) Error bars represent SEM. * p < 0.05 (Independent t-test)

The activity of GalR1-expressing neurons in the two areas showed a highly distinct relationship to behavioral events. For example, the vPFC neurons showed an increase in activity just prior to trial initiation (Fig. 4c, e). On trials that were completed correctly, the activity remained elevated including during cue presentation, suggesting a close relationship with the animal’s attention. However, the corresponding vHC neurons did not show such elevation but remained low throughout the same period. During incorrect trials, the same pattern in the vPFC and vHC was observed but the activity levels were lower compared with correct trials (Fig. 4d). Later, after the response, the vPFC signal increased presumably in anticipation of reward, but only on correct trials (Fig. 4d, f). In contrast, in trials with omissions or in the timeout following premature responses (Fig. 4g-j), the vHC neurons showed elevated activity, suggesting that these GalR1 expressing neurons may signal cognitive errors or negative events. Together, these data suggest that the GalR1-expressing neurons in the vPFC and vHC have highly distinct activity profiles that are linked to unique cognitive signals and behavioral outcomes.

### Activity levels in vPFC GalR1-expressing neurons predict behavioral outcome

To confirm the relationship between the activity levels of GalR1-positive neurons and the behavioral response, we compared the level of activity for each trial outcome: 1) before the rat initiated the trial, 2) during the pre-stimulus interval, and 3) when the cue was presented for each brain region over 1 sec time bins (Fig. 5). In the vPFC, the highest level of activity during the pre-stimulus interval was associated with a future correct response, while lower activity predicted inappropriate behavior namely incorrect responses, premature responses, or omissions (Fig. 5a). In the vHC, calcium activity was higher during the pre-stimulus interval when the animals later made an omission, but no statistical differences were observed (Fig. 5b). Thus, the activity of GalR1-expressing neurons while animals perform the 5-choice task are both region- and response-specific. These results are in line with the idea that the level of prefrontal neuronal activity correlates with behavioral outcome [29, 30].

**Figure 5.**
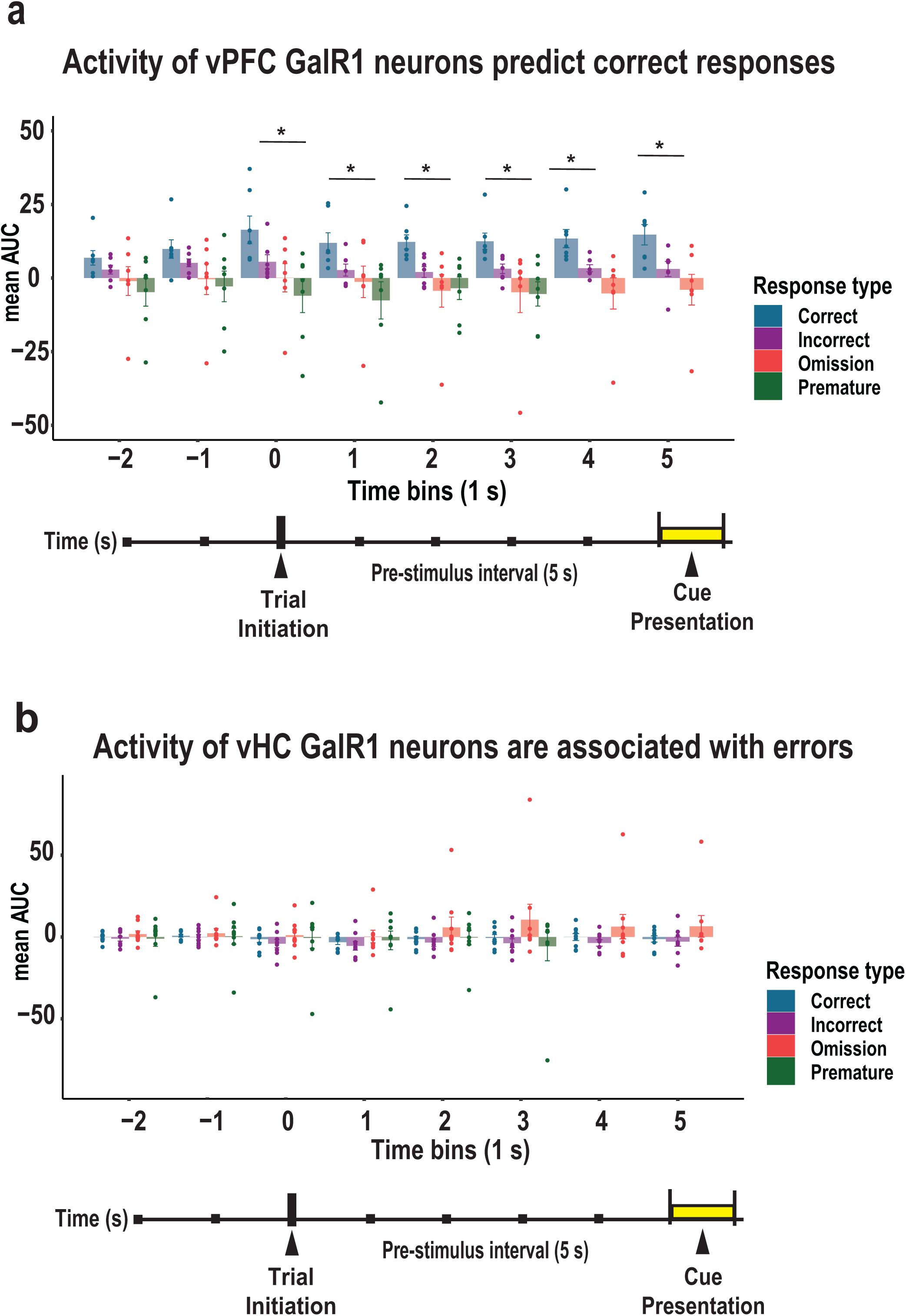
Activity of vPFC and vHC GalR1-expressing neurons predict behavioral outcomes (a) Comparison of activity in vPFC GalR1-expressing neurons expressed as area under the curve (AUC) for each response outcome. Time starts 2 seconds before trial initiation until the end of the cue presentation divided into 1 second time bins (color coded by response type). Bars represent mean AUC and error bars represent SEM. * p < 0.05 (One-way ANOVA). No changes were observed before trial initiation [2 sec before F(3, 24) = 1.826, p = 0.169; 1 sec before F(3,24) = 1.968, p = 0.146]. When the trial was initiated, and for 1 sec bins thereafter, the activity of vPFC GalR1was associated with a future correct response: trial initiation (F(3, 24) = 4.328, p = 0.014. Post hoc comparison: p = 0.355 for correct vs incorrect, p = 0.082 for correct vs omission, p = 0.01 for correct vs premature, p = 0.835 for incorrect vs omission, p = 0.305 for incorrect vs premature, p = 0.782 for omission vs premature ; 1 second after trial initiation: F(3, 24) = 3.154, p = 0.043, Post hoc comparison: p = 0.502 for correct vs incorrect, p = 0.205 for correct vs omission, p = 0.03 for correct vs premature, p = 0.926 for incorrect vs omission, p = 0.408 for incorrect vs premature, p = 0.77 for omission vs premature; 2 seconds after trial initiation: F(3, 24) = 4.274, p = 0.015, Post hoc comparison: p = 0.232 for correct vs incorrect, p = 0.02 for correct vs omission, p = 0.029 for correct vs premature, p = 0.618 for incorrect vs omission p = 0.719 for incorrect vs premature, p = 0.998 for omission vs premature; 3 seconds after trial onset: F(3, 24) = 3.731, p = 0.025, Post hoc comparison: p = 0.44 for correct vs incorrect, p = 0.044 for correct vs omission, p = 0.036 for correct vs premature, p = 0.574 for incorrect vs omission p = 0.516 for incorrect vs premature, p = 1.0 for omission vs premature; 4 seconds after trial onset: F(2, 24) = 6.649, p = 0.007, Post hoc comparison: p = 0.148 for correct vs incorrect, p = 0.005 for correct vs omission, p = 0.148 for incorrect vs omission; cue: F(2, 24) = 5.838, p = 0.011, Post hoc comparison: p = 0.116 for correct vs incorrect, p = 0.009 for correct vs omission, p = 0.427 for incorrect vs omission]. **(b)** Comparison of activity in vHC GalR1-expressing neurons expressed as the area under the curve (AUC) of each response outcome. Time starts 2 seconds before trial initiation until the end of the cue presentation divided into 1 second time bins (color coded by response type). Bars represent mean AUC and error bars represent SEM. (One-way ANOVA). No differences were observed before or after trial initiation [- 2 seconds: F(3,24) = 0.243, p = 0.866; -1 second: F93, 24) = 0.115, p = 0.95; trial initiation: F(3, 24) = 0.326, p = 0.807; 1 second: F(3, 24) = 0.402, p = 0.753; 2 seconds: F(3, 24) = 0.907, p = 0.449; 3 seconds: F(3, 24) = 1.181, p = 0.332; 4 seconds: F(2, 24) = 1.202, p = 0.318; cue: F(2, 24) =1.35, p = 0.278].

## Discussion

In the present study, we used multiple approaches to characterize the functional differences between GalR1-expressing neurons of the vPFC and the vHC and their involvement in mechanisms of attentional control. There were several findings: 1) GalR1 was predominantly expressed in glutamatergic neurons in both regions, 2) optical activation of GalR1-expressing neurons in the vPFC, but not vHC, influenced selective attention and impulse control, 3) the activity of vPFC neurons predicted successful response outcomes, and 4) the activity of vHC neurons were associated with inappropriate responses. Together, these data provide evidence that GalR1 expressing neurons produce region- and response-specific intrinsic activity in the vPFC and vHC to influence the expression of cognitive-executive behaviors.

### GalR1 distribution in the vPFC and vHC

The presence of GalR1 in the vPFC and vHC has been previously described [14, 15, 26]. We now add to this information that these receptors are mostly expressed in layer 5 of the vPFC and the pyramidal layers of the vHC (particularly the vCA1 and vSub), which are the main output layers in both regions [31, 32]. The receptors’ locations indicate the possibility for galanin to influence the actions of these neurons on target areas. While those layers are comprised of diverse subclasses of neurons, we found that GalR1 was expressed predominantly in putative pyramidal neurons expressing VGLUT1. It bears mentioning that a small proportion of GalR1-expressing neurons in both areas were identified as GABAergic, which may further shape the influence of galanin over PFC or hippocampal circuits.

With *in situ* hybridization, we were able to visualize mRNA but not the active protein and its ultimate subcellular location. We assume, therefore, that the majority of GalR1 receptors are located on the soma, but receptors can also be present on dendritic trees where they could modulate the integration of signal coming from specific inputs. Although the expression of GalR1 has been detected on proximal dendrites with immunostaining [33], the specificity of the antibodies targeting GalR1 has been questioned [34]. In addition, GalR1 may be located on pre-synaptic terminals [35]. The multiple possibilities for GalR1 cell-type and subcellular distribution suggest that galanin modulation of neural circuits is complex and involves diverse mechanisms within each region. The exact subcellular expression of the galanin receptor remains to be determined before its contribution to the microcircuit’s dynamics can be assessed. Our data revealed a similar distribution of GalR1 expression in the vPFC and vHC and suggests that the region-specificity of galanin modulation of executive functions might be due to the intrinsic differences between prefrontal and hippocampal neuronal populations and their respective contributions to behavior.

### GalR1-expressing neurons of the vPFC contribute to attentional control

The optical manipulation of GalR1-expressing neurons demonstrated strong differences in the involvement of vPFC and vHC neurons during performance of the 5-choice task. The data demonstrated that GalR1-expressing neurons in the vPFC are directly involved in the control of attention since activating these neurons specifically decreased the animals’ ability to accurately detect the target while increasing omissions and decreasing impulsive premature responses. These data are consistent with lesion and pharmacological studies that have implicated an important role for the vPFC in the normal control of impulsive urges [27, 36–38]. In contrast to lesions or drug infusions, optical perturbation affords temporally precise causal manipulations. In our case, the stimulation was selectively applied during the 5 s of the pre-stimulus interval and in only half of the total trials of a session. Opto-inhibition of vPFC neurons during the entire pre-stimulus interval has previously been shown to increase accuracy in the 5-choice task, whereas distracting the animal by inhibiting the neurons two seconds before the cue presentation, has the opposite effect [39]. Since the GalR1-expressing neurons represent only a portion of the vPFC and vHC neurons it is unlikely that the stimulation captured the full effects observed by manipulations affecting the entire region.

In contrast, stimulation of GalR1-expressing neurons in the vHC resulted in an increase in omissions only. Although this finding was similar to the effects of vPFC stimulation (see Fig. 2f, j), it was in marked contrast to the more global effects of hippocampal disinhibition previously shown to induce attentional deficits [40]. Moreover, rats with HC lesions display exaggerated and persistent responding; have long lasting increases in premature responses [41], are resistant to extinction [42, 43] and display reward induced stereotypy [44]. One parsimonious explanation is that the selective targeting of vHC GalR1 neurons are more subtle in their effects relative to large global vHC lesions. A more speculative hypothesis is that an intact vHC facilitates the evaluation of negative feedback following inappropriate actions to adapt choices accordingly. In humans, hippocampal signals have been shown to differentiate between positive and negative feedback [45, 46]. It is feasible therefore that a lesioned vHC would diminish the monitoring of such feedback so that unfavorable consequences of timeout/no reward could ostensibly lead to repeated errors. Finally, GalR1-expressing neurons represent a portion of the vPFC and vHC neurons and it is likely that their stimulation will not recapitulate the effects observed by manipulations affecting almost the entire region. Nevertheless, while the two regions contribute differentially, the vPFC is a stronger driver of behavior in the 5-choice task.

### Activity of vPFC and vHC GalR1-expressing neurons predict behavioral outcomes

The photometry traces of calcium activity shed light on the intrinsic activity of GalR1- expressing neurons during task performance. One major finding was the region-specific activity patterns for each behavioral outcome. In the vPFC, GalR1 expressing neurons that predicted a correct response sustained a high level of activity during the entire pre-stimulus interval, whereas incorrect responses including premature responses were associated with a lower magnitude of response [28, 47]. This was in contrast with trials which led to an omission for which the GalR1 vPFC neurons showed no change in activity. Thus, the activity of vPFC GalR1-expressing neurons may be an indicator of the level of task engagement with high levels of sustained activity reflecting full engagement resulting in successful goal directed behavior [48]. Studies in rats, monkeys, and humans [49, 27, 50] consistently identify the PFC as the key site for attentional processing and the driver of coordinated goal directed actions, but the link between intrinsic region-specific PFC activity and behavioral outcome is largely missing. Our data confirm that recruitment of vPFC GalR1-expressing neurons is critical for sustained attentional processing.

The activity patterns for the vHC GalR1-expressing neurons were more nuanced. Overall, the signal in vHC neurons was lower than vPFC neurons during both correct and incorrect trials. Thus, vHC GalR1-expressing neurons may not participate in attentional mechanisms that predict successful outcomes. Instead, the vHC signal was more associated with errors or inappropriate responses during which the activity of vHC neurons became higher than the vPFC. The elevated activity preceding an omission, for example, is consistent with the increased omissions observed following optogenetic stimulation. Intriguingly, the activity of vHC GalR1 neurons elevated substantially during the timeout period immediately following an impulsive premature response. Since timeouts provide negative feedback (i.e., no reward), it is conceivable that the heightened activity in the vHC neurons represents an emotional state of disappointment or frustration. There is much evidence that the vHC is intimately tied to negative emotional states [51, 52]. One possibility is that GalR1-expressing neurons in the vHC may inappropriately enhance attention towards negative events thereby promoting behaviors with strong affective components such as anxiety and depression [53, 54]. This hypothesis needs to be tested directly.

In conclusion, our findings suggest an important role for GalR1-expressing neurons of the vPFC and vHC in attentional processing. Neurons in the vPFC and vHC differentially signal cognitive mechanisms of attention and impulse control, which explains why pharmacological activation of GalR1, which hyperpolarizes neurons [55], leads to opposing effects on the attentional control of behavior [25]. It also suggests that synchronously activating a population of neurons can reveal very different patterns of behavior than manipulating a specific receptor with pharmacology. Although the vHC and vPFC interact both anatomically and functionally, our data highlight the fundamental differences between these structures in executive control behaviors. How GalR1-positive neurons respond differently than GalR1-negative neurons in vPFC and vHC structures requires further investigation.

## Methods and Materials

### Animals

A total of 89 adult male Long-Evans rats (Envigo, Indianapolis, IN, USA) were housed in pairs in a temperature-controlled room (23.3 °C) under a 12 h light/dark cycle. About two weeks after their arrival, animals were food restricted and maintained at 85% of their free-feeding weight throughout the experiments. All experimental procedures were approved by NIMH Institutional Animal Care and Use Committee, in accordance with the NIH guidelines for the use of animals.

### Histology and immunohistochemistry

For immunofluorescence staining, naïve animals were perfused transcardially with a working solution of PBS (1X) followed by 4% paraformaldehyde in phosphate-buffered saline. The brains were extracted and postfixed in 4% paraformaldehyde. After dehydration by immersion in 25% sucrose, the brains were cryo-sectioned at 40 μm thickness. Galanin fibers were labeled using rabbit anti-galanin primary antibody (Thermo Fisher Scientific #PA5-6209, 1:500) and Alexa 647 goat anti-rabbit secondary antibody (Thermo Fisher Scientific #A27040, 1:500). Sections were mounted onto slides and coverslipped with the VectaShield HardSet Antifade mounting medium with DAPI (Vector Laboratories H-1500-10). Images were acquired with a Zeiss Axioscan at 10x magnification. For quantification of galanin fibers in the respective regions of interest, we examined 3-4 sections for each animal from each region. All image analysis was performed with ImageJ (NIH, Bethesda, MA, USA, https://imagej.nih.gov/ij/download.html). We first split the composite image of the section into two channels to create a gray-scale image (Fig S1). The green channel (galanin) images were then converted to reduce background and increase visibility of fibers using the FeatureJ: Hessian plugin in image J with the smallest eigen value and a smoothing scale of 1.0. Contrast was enhanced by 0.01%. With the created ROIs, the means were recorded. Data was reported as a mean for each section.

### RNAscope in situ-hybridization (ISH)

We applied RNAscope ISH to detect the expression of GalR1, Slc17a7 (VGluT1), Slc32a1 (VGAT) and tdTomato mRNA in the vPFC and vHC using the RNAscope Fluorescent Multiplex Assay (Advanced Cell Diagnostics, Newark, CA, USA). We mounted 16 µm sections from flash-frozen brains directly onto Superfrost Plus slides (Thermo Fisher Scientific, Waltham, MA, USA). Briefly, sections were fixated with cold 4% PFA in PBS for 1h, washed with PBS, and dehydrated in a series of increasing concentrations of ethanol in water (50%, 70%, and twice 100%). The sections were then incubated with Protease IV for 30 min and washed in distilled water. Subsequently, probes of interest were applied to the sections, and hybridization was carried on for 2h at 40 ℃. This was followed by 4 amplification steps for 30 min, 15min, 30 min and 15 min respectively at 40 ℃. Each amplification step was followed by two washes in wash buffer. Sections were finally cover slipped with mounting medium containing DAPI. Images were acquired using a Leica Stellaris confocal microscope (Leica Microsystems, Wetzlar, Germany) at 40x magnification or on a Zeiss Axioscan (Zeiss, Oberkochen, Germany) at 20x magnification. Images were further processed in ImageJ, and Cell Profiler software (Broad Institute, Cambridge, MA, USA, https://cellprofiler.org) provided quantification of the expression of the mRNAs of interest.

### Viruses

Adeno-associated virus (serotype 1) expressing Cre recombinase under the promoter of the galanin receptor 1 (GalR1) was produced by the Viral Vector Core, National Institute of Neurological Disorders and Stroke (Bethesda, MD, USA) (titer 3×10¹² vg/mL). The following viruses were purchased from Addgene (Watertown. MA, USA): AAV1-EF1a-Flex-hChR2(H134)- EYFP-WPRE-HGHpA (Addgene viral prep # 20298-AAV1, titer 7×10¹² vg/mL, gift from Karl Deisseroth), AAV1 CAG-LSL-tdTomato (Addgene viral prep #100048-AAV1, gift from Hongkui Zeng), AAV1-Flex-tdTomato (Addgene viral prep # 28306-AAV1, titer 1×10¹³ vg/mL, gift from Edward Boyden), AAV1-CAG-Flex-EGFP-WPRE (Addgene viral prep # 51502-AAV1, titer 1×10¹³ vg/mL, gift from Hongkui Zeng), and AAV1-CAG-Flex-jGCaMP7f-WPRE (Addgene viral prep # 104496-AAV1, titer 7×10¹² vg/mL, gift from Douglas Kim & GENIE Project).

### Viral injections

For all procedures involving local injections of virus, rats were anaesthetized with isoflurane gas (5% induction, 2% maintenance) and placed in a stereotaxic frame fitted with atraumatic ear bars (David Kopf Instruments, Tujanga, CA, USA). The scalp was retracted to expose the skull, and craniotomies were made directly above the target brain regions. The following coordinates were used for injections: vPFC (AP +3.24, ML 0.6, DV -3.7 from dura), vHC (AP -5.0, ML 5.4, DV -6.7 from dura). The same coordinates were used for these regions for each project.

### Validation of galanin receptor 1-Cre virus

A cocktail of AAV1-GalR1-Cre with AAV1 Cre-dependent expressing tdTomato was injected directly into the vPFC (AP +3.24, ML 0.6, DV -3.7). The fourth cerebellum lobule (4Cb) (AP - 9.72, ML 1.9, DV -1.8) was also injected as a control area that lacks GalR1. All DV readings were taken from dura. For all injections, a total of 0.1 - 0.3 µl was infused at a rate of 0.1 µl / min.

### Anatomical projections of GalR1-expressing neurons

To map the projections of vPFC and vHC GalR1-positive neurons, the same animals were also injected with cocktails of AAV1-GalR1-Cre with Cre-dependent AAV1 expressing GFP in the vPFC or tdTomato in the vHC. For all injections, a total of 0.1 - 0.3 µl was infused at a rate of µl / min.

### Behavioral procedure: 5-choice task

Two weeks following stereotaxic placement of fiber implants (see below), rats were trained to accurately detect the occurrence of a brief visual target (a white square) in the 5-choice attentional task using the touchscreen operant platform. Full details of the apparatus and behavioral procedure can be found in Messanvi et al. (2020) [25]. Some adaptations to the apparatus were necessary to enable the optogenetic and fiber photometry settings. In brief, animals were first habituated to moving around freely in the operant chamber while tethered to the patch cord (Doric Lenses). The patch cord was connected to the fiber-optic rotary joint (Doric Lenses) thereby allowing the animals to move freely inside the chambers. While tethered the animals were pretrained to: a) successfully enter the food magazine, b) reliably touch the screen with their nose, collect food reward (Dustless Precision Pellets, Bio-serv, Flemington NJ, USA), and d) initiate trials.

The patch cord was disconnected when the rats were trained for the main task (∼20 days). A daily session consisted of 100 completed trials or was terminated after 35 min, whichever came first. Rats initiated the trial by making a nose entry into an illuminated food magazine. Following a 5 s interval, a brief white square stimulus was presented pseudo-randomly in one of five spatial locations on the screen. Animals made a response on the screen by touching the stimulus with its nose. A correct detection within 5 s was rewarded with a single food pellet. Following a 2 s food consumption time, the next trial was signaled by the illumination of the food magazine. Incorrect responses, failure to respond (omissions) or responses ‘before’ the stimulus presentation (premature response) terminated the trial during which the chamber was illuminated for 5 s and reward was not delivered. In addition, latencies to initiate each trial, make a response, and collect reward were recorded.

In each session, the visual target was presented an equal number of times in one of five locations in a pseudo-random order. During training, the target duration and response window were set at 10 s. These variables were reduced on subsequent sessions according to the individual animal’s performance until the target duration was 1 s and the response time was 5 s. These served as the baseline parameters. When rats displayed greater than 75% accuracy with less than 30% omissions at the baseline parameters, they were ready for optical stimulation.

The apparatus and online data collection for each chamber were controlled by a Dell computer connected to an Animal Behavior Environmental Test (ABET) software (Lafayette Instruments Company, Lafayette, IN, USA) interfaced with the Whisker control system for research [56].

### Optogenetic stimulation

We targeted the vPFC and vHC in separate groups of animals. A cocktail of AAV1-GalR1- Cre with Cre-dependent AAV1 expressing ChR2, or tdTomato as a control, was injected bilaterally into the vPFC or the vHC (200 nl). Subsequently, 0.3 mm dorsal to the viral injection, dual fiber-optic cannulas were implanted in the vPFC (200 μm core diameter, 0.37 NA, 6mm length; Doric Lenses, Quebec, QC, Canada). Bilateral fiber-optic cannulas were implanted in the vHC (200 μm core diameter, 0.39 NA, 8mm length; Thorlabs).

Once rats had acquired the baseline parameters of the 5-choice task, they were re-habituated to the patch cord (∼3 days) and remained tethered to the patch cord during the remaining test sessions. Optical stimulation (473 nm, 4 mW intensity at the end of the dual optic fiber tip, 5 ms pulse duration, at 40Hz) was delivered using a laser system (LRS-0473, Laserglow Technologies, North York, ON, Canada) for 5 s for the entire duration of the pre-stimulus interval. Half of the trials were stimulated (ON trials) and the other half were not (OFF trials). Stimulated trials were distributed pseudo-randomly throughout the session, across the five locations.

### Fiber photometry

We injected AAV1-GalR1-Cre and AAV1-CAG-Flex-jGCaMP7f-WPRE (200 nl) viruses into the vPFC and vHC. Fiber optic cannulas (400 μm core diameter, 0.66 NA, 5 or 8mm length for vPFC and vHC respectively; Doric Lenses) were placed 0.1 mm dorsal to the viral injection. In all cases, cannulas were affixed with dental cement and, stainless steel screws to secure them in place. Two weeks after surgery, the rats were trained on the 5-choice task until stable baseline performance (∼ 20 days).

Fiber photometry data were acquired with the RZ10X processor integrated with Synapse Software v.96 (Tucker-Davis Technologies, Alachua, FL, USA). Lights emitted from LEDs (465 nm modulated at 330 Hz to excite GCamP7f, and 405 nm modulated at 210 Hz for the isosbestic control) were relayed to the mini cubes (Doric Lenses) via attenuator patch cords. Lights were then conveyed to the fiber-optic cannulas implanted in the rats’ brain, via a pigtailed rotary joint (Doric Lenses) and two low-autofluorescence optic fibers (400 μm core diameter, NA 0.48, Doric Lenses). The signals from the brain were sent back to the mini cubes for filtration, detected by the photosensors, and finally demodulated in the Synapse software. In parallel, time stamps of the behavioral events (initiation, cue, response types, reward collection) from ABET were sent to the fiber photometry system through a TTL breakout adapter (Lafayette Instruments).

Raw fluorescence signals and time stamps for signals and behavioral data were extracted by importing the TDT files into the fiber photometry Modular Analysis Tool (pMAT) [57]. Extracted data were further processed using a custom-written R code, to separate the signal around specific events. We used again pMAT to calculate the Z-score and area under the curve (AUC) values. The 2 s preceding trial initiation were used as a baseline to generate normalized Z-score values. The AUC values preceding and following specific events were averaged over specific time bins (the durations of the different time bins are specified in the figures legends). Data were calculated for each trial, then averaged over the session, and finally over experimental groups.

### Verification of fiber placement and viral expression

Animals were transcardially perfused with a working solution of PBS (1X) followed by 4% paraformaldehyde in phosphate-buffered saline. The brains were extracted and postfixed in 4% paraformaldehyde. After dehydration by immersion in 25% sucrose, the brains were cryo-sectioned at 40 μm thickness. Every other section was mounted on glass slides and cover-slipped with mounting medium containing DAPI (Vector Laboratories, Newark, CA, USA) for fluorescence microscopic imaging. Pictures were taken using an Axioscan Z1 (Zeiss) at 10x magnification, and animals with misplaced cannulas or viral expression were excluded from the analysis.

### Statistical analysis

Statistical analyses were performed using SPSS (29.0.1.0, IBM, Armonk, NY, USA). For the optogenetics experiments, the effects of laser stimulation, brain regions and their interaction were determined using a mixed ANOVA with repeated measures. When interaction of the two factors was found to be significant, post-hoc pairwise comparisons (with Bonferroni correction) were performed. For the photometry experiments, comparison of the signals between brain regions was performed with independent t-tests. Comparison of signals between behavioral outcomes was performed using a one-way ANOVA, and post-hoc pairwise comparisons (with Bonferroni correction) were performed when F ratios were significant. The criterion for significance for all analyses was set at p < 0.05. Data are reported as mean ± SEM.

## Acknowledgments

This research was supported by the Intramural Research Program of the National Institute of Mental Health (ZIA MH002951 and ZIC MH002952 to YC). We thank Raymond Fields from the NINDS Viral Vector Core for creating the GalR1-Cre construct, the NIMH Systems Neuroscience Imaging Resource (SNIR) for their technical assistance with imaging, and the NIMH Rodent Behavioral Core (RBC) for their assistance with surgery and behavioral experiments. CS is now at UMass Chan Medical School, KB is now at the Vienna Biocenter in Austria, and FM is now faculty at American University, Washington DC.

## Data availability

All relevant data will be deposited in Mendeley Data public repository.

---

**Supplementary Figure 1.**
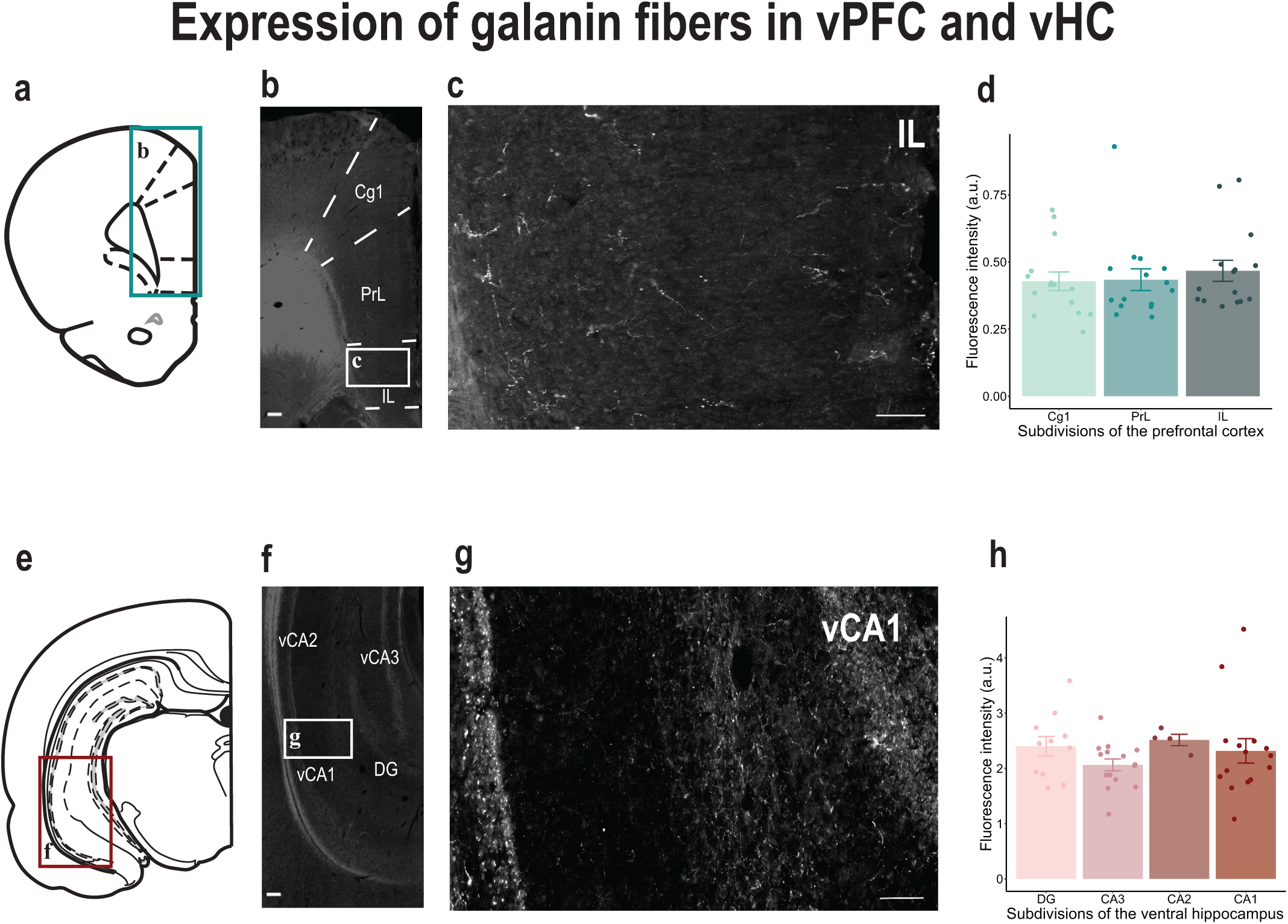
Expression of galanin fibers in the vPFC and the vHC. **(a)** Schematic representing the location of the area analyzed in the PFC. **(b)** Microphotograph shows the distribution of galanin fibers and terminals in the PFC subdivisions (scale bar: 200 µm). **(c)** Magnified image of the galanin-containing fibers and terminals in the IL (scale bar: 200 µm). **(d)** Quantification of the mean density of galanin immunofluorescence in the three PFC subdivisions (Cg1: 0.43 ± 0.035, PrL: 0.43 ± 0.04, IL: 0.47 ± 0.04; au: arbitrary unit; N = 5 animals, 15 sections per region). Bar chart represents mean ± SEM. Dots represent individual animals. **(e)** Schematic representing the location of the area analyzed in the HC. **(f)** Microphotograph shows the distribution of galanin fibers and terminals in the vHC subdivisions (scale bar: 200 µm). **(g)** Magnified image of the galanin-containing fibers and terminals in the vCA1 (scale bar: 200 µm). **(h)** Quantification of the mean density of galanin immunofluorescence in the vHC subdivisions (CA1: 2.32 ± 0.22, CA2: 2;51 ± 0.10, CA3: 2.06 ± 0.11, DG: 2.40 ± 0.18; au: arbitrary unit; N = 1 animal, 4 to 15 sections per region). Bar chart represents mean ± SEM. Dots represent individual animals.

**Supplementary Figure 2.**
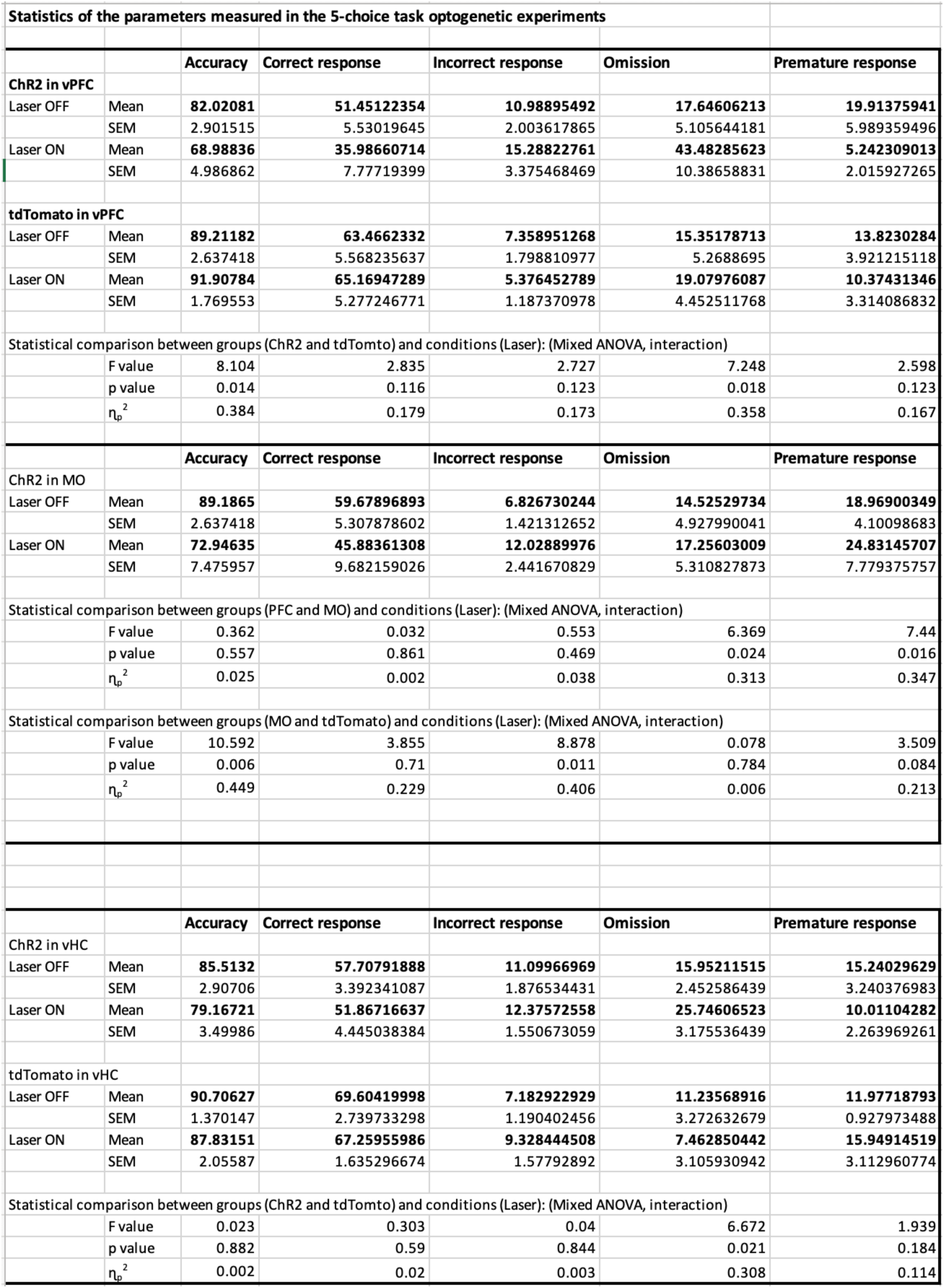
**Statistics for the main behavioral outcomes assessed in the 5-choice task for the optogenetic experiment.**

**Supplementary Figure 3.**
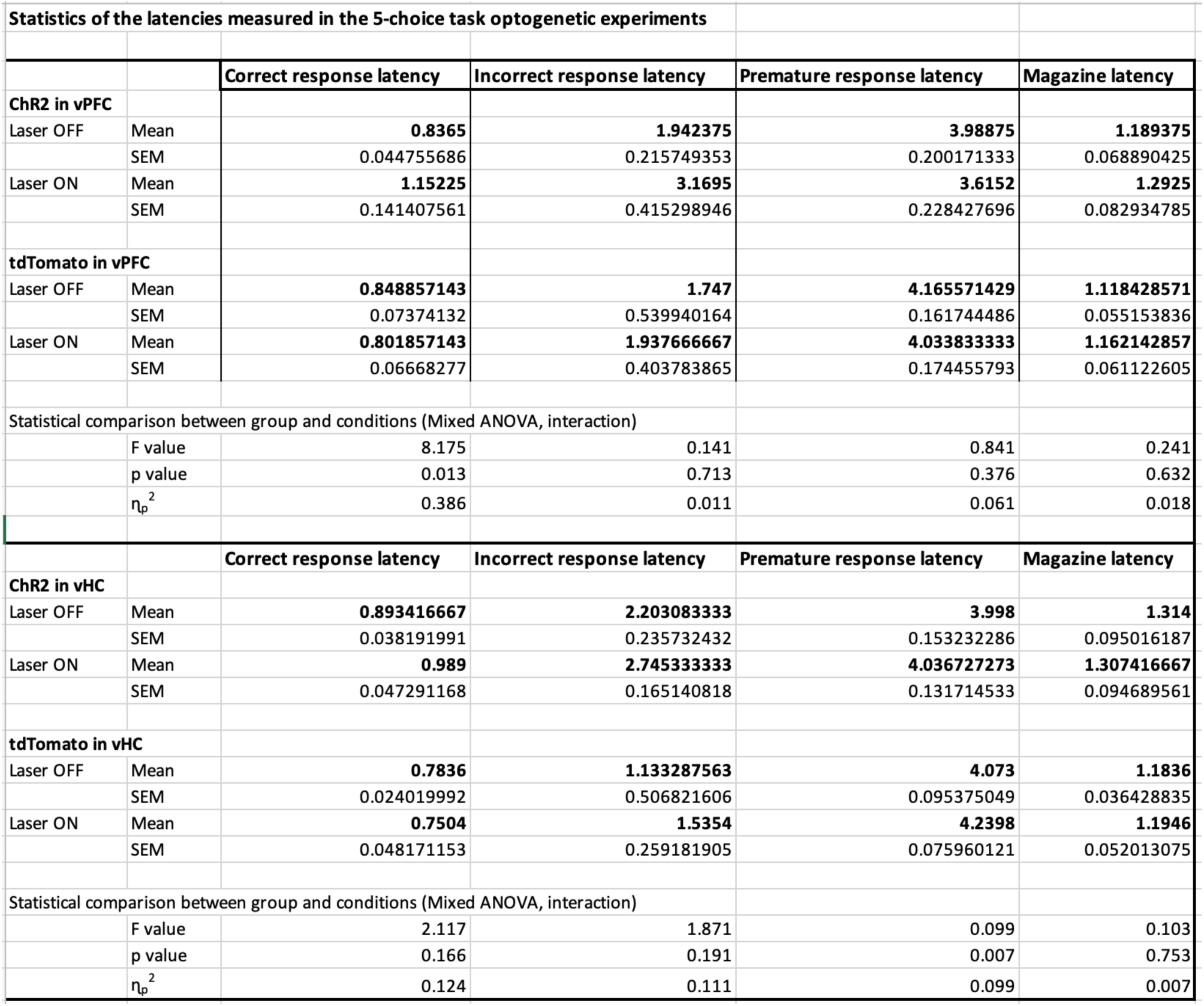
**Statistics for the latency measures in the 5-choice task for the optogenetic experiment.**

**Supplementary Figure 4.**
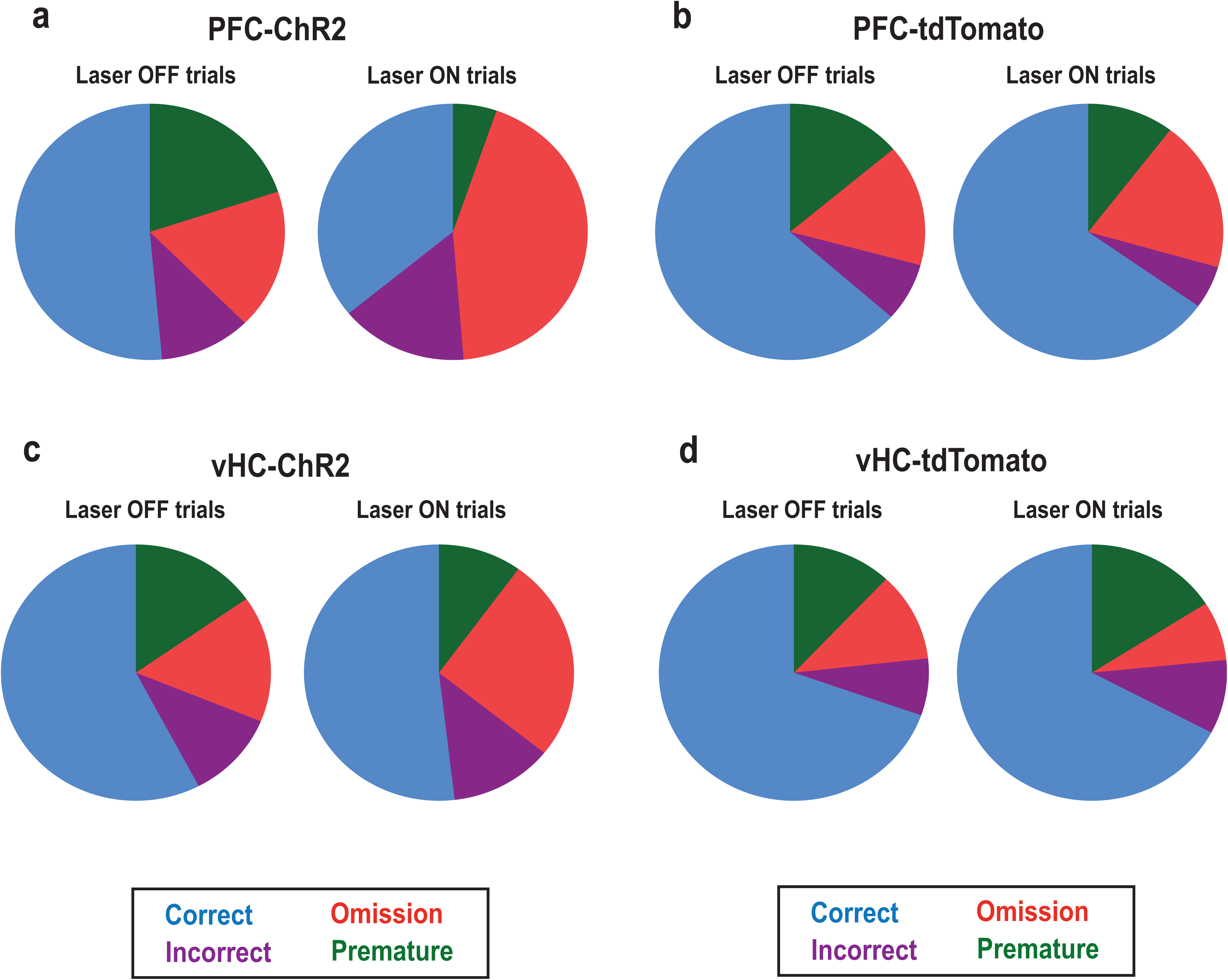
Proportion of response types between laser conditions. **(a)** Proportion of response types in non-stimulated and stimulated trials for vPFC-ChR2 animals (rats n = 8). **(b)** Proportion of response types in non-stimulated and stimulated trials for vPFC- tdTomato animals (rats n = 7). **(c)** Proportion of response types in non-stimulated and stimulated trials for vHC-ChR2 animals (rats n = 12). **(d)** Proportion of response types in non-stimulated and stimulated trials for vHC-tdTomato animals (rats n = 5).

**Supplementary Figure 5.**
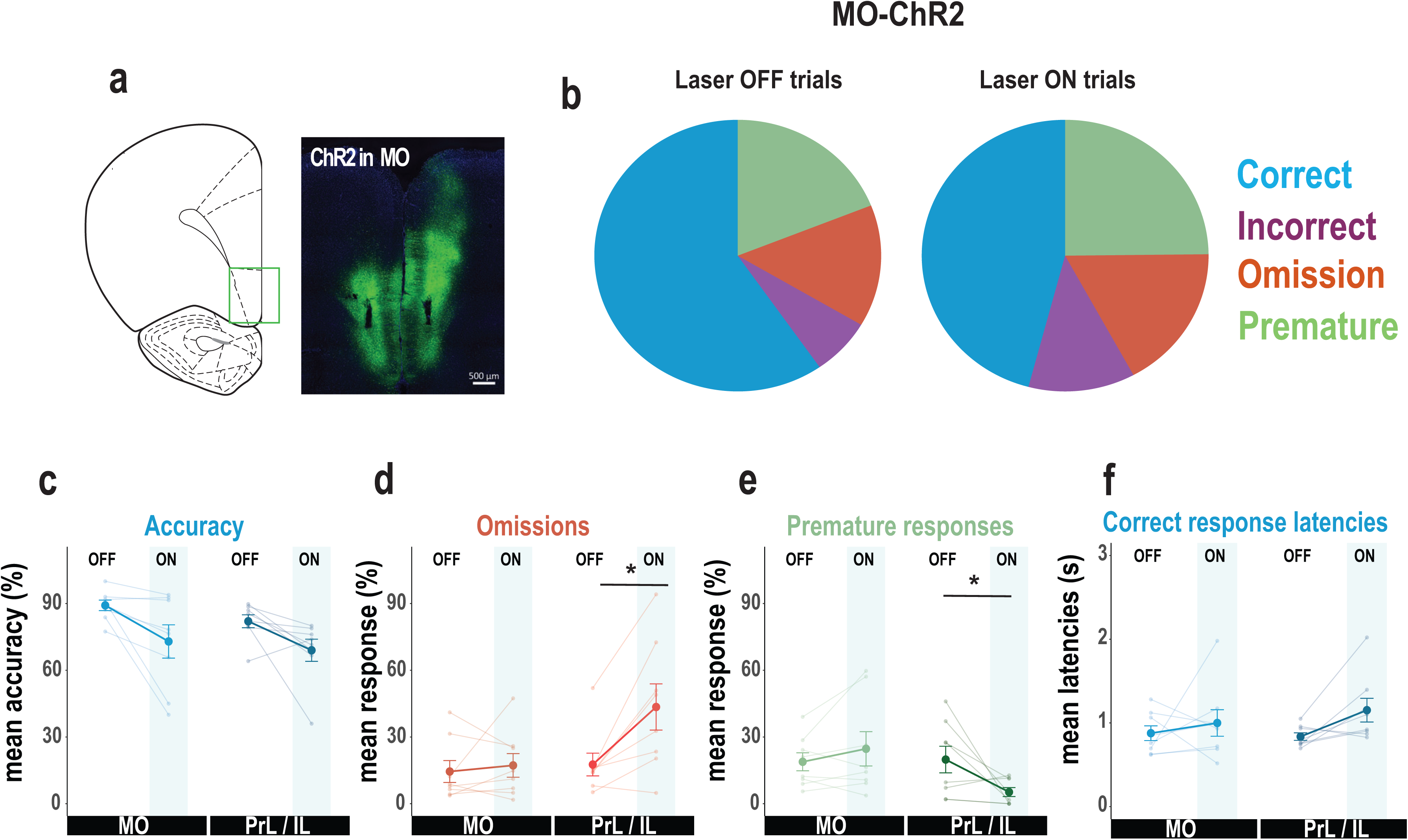
Comparison of MO and PrL/IL optical stimulation on behavioral responses. **(a)** Schematic of brain section showing location of viral expression in the medial orbital region (MO) with corresponding photomicrograph. **(b)** Proportion of response types in non-stimulated and stimulated trials for animals with optic fiber in the MO brain region (rats n = 8). **(c-f)** Behavioral effects of optical stimulation for MO or PL/IL GalR1-expressing cells on accuracy, omissions, premature responses, and correct response latencies (MO: n = 8; IL: n= 8). Error bars represent SEM. * p < 0.05 (paired t-test between conditions after significant interaction effect in mixed ANOVA)

**Supplementary Figure 6.**
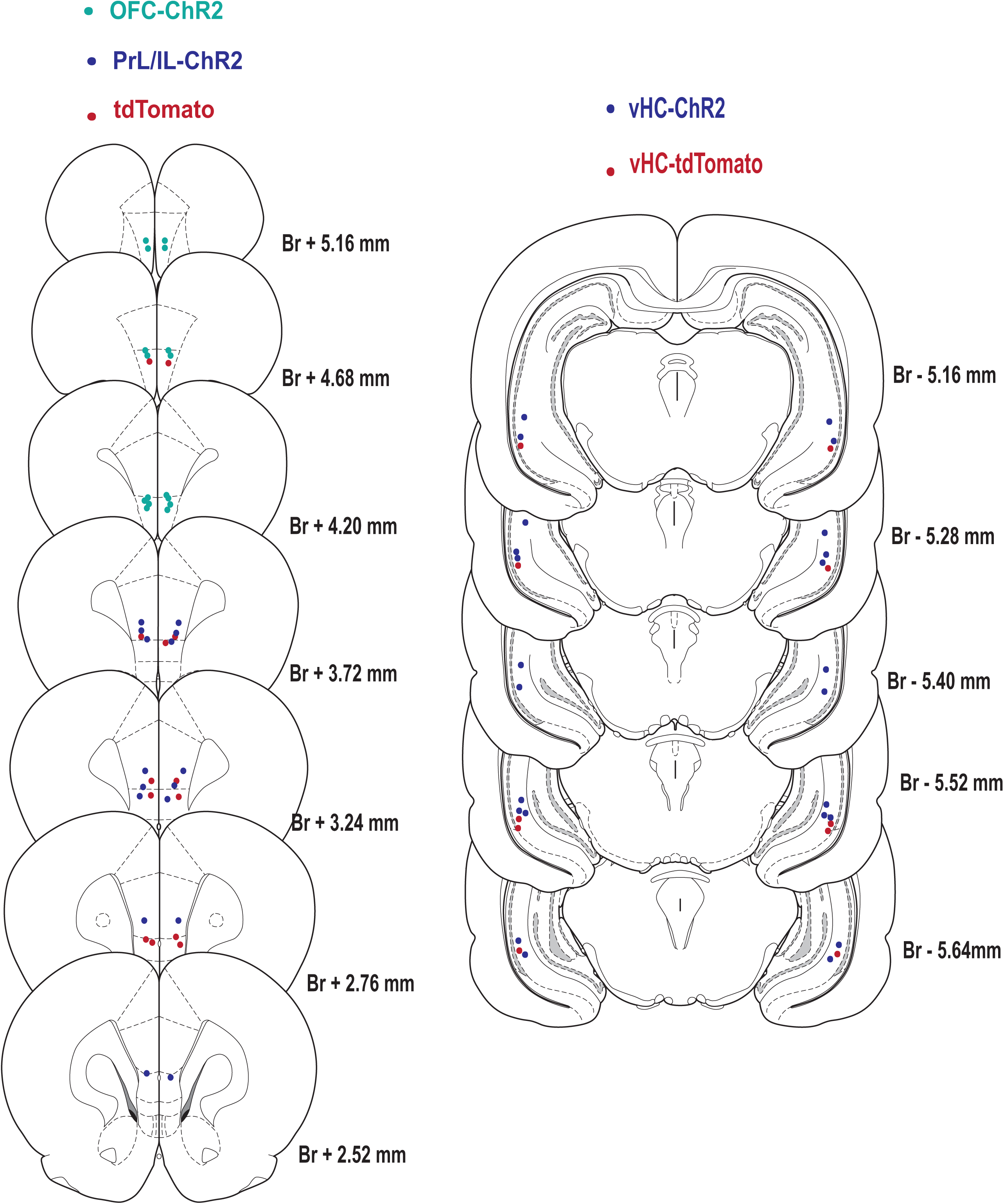
Location of optical fibers in the vPFC and vHC for the optogenetics experiments.

**Supplementary Figure 7.**
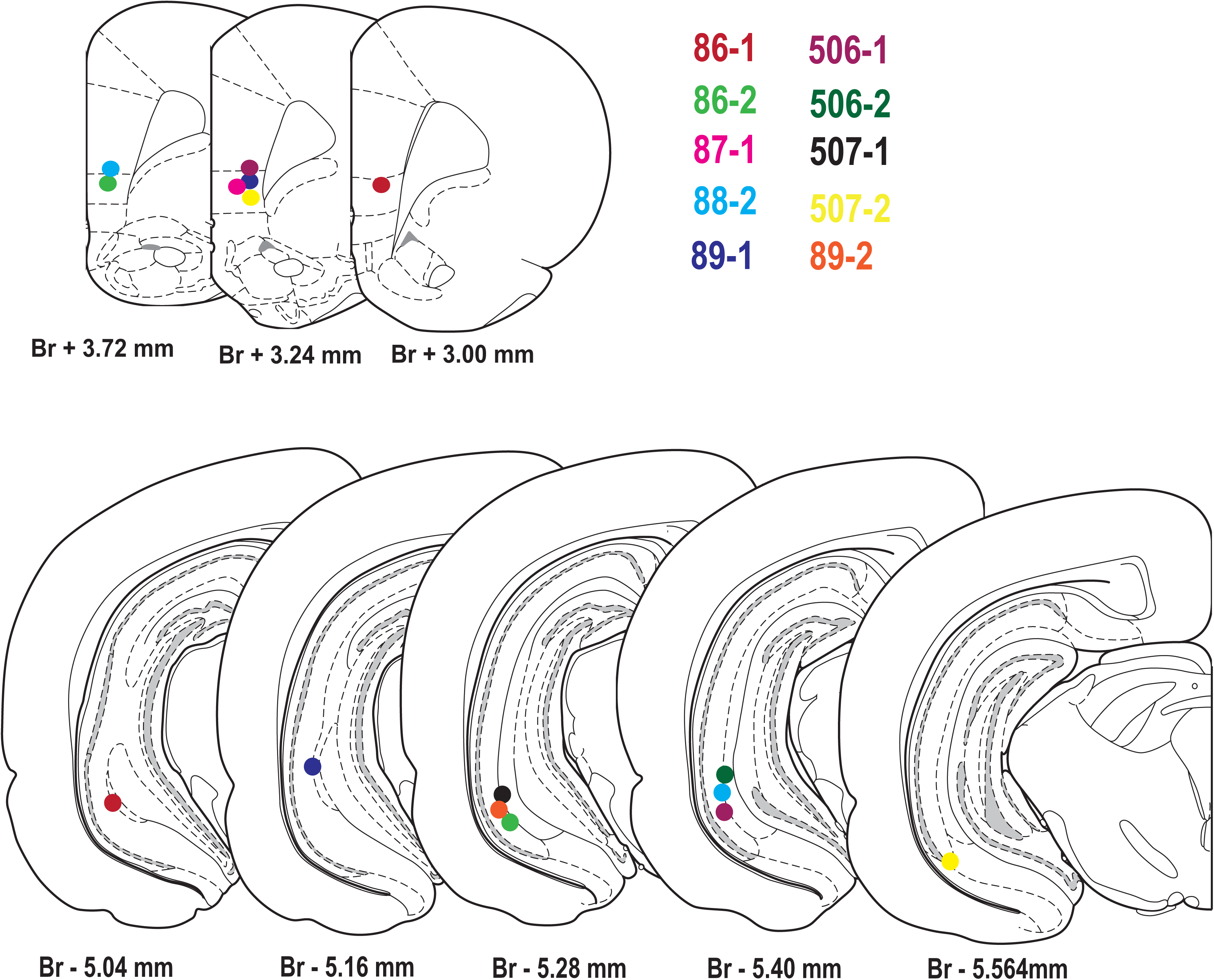
**Location of optical fibers in the vPFC and vHC for the photometry experiments.**

